# Optimizing PDIA1 Inhibition as a Strategy to Inhibit NLRP3 Inflammasome Activation and Activity

**DOI:** 10.1101/2025.03.21.644355

**Authors:** Chavin Buasakdi, Caroline R. Stanton, Prerona Bora, Priyadarshini Chatterjee, Michael J. Bollong, R. Luke Wiseman

**Author notes:** To whom correspondence should be addressed R. Luke Wiseman, Department of Molecular and Cellular Biology, The Scripps Research Institute, La Jolla, CA 92037, Michael J. Bollong, Department of Chemistry, The Scripps Research Institute, La Jolla, CA 92037.

## Abstract

The NLRP3 inflammasome is a protein complex that promotes pro-inflammatory signaling as part of the innate immune response. Hyperactivation of the NLRP3 inflammasome has been implicated in many inflammatory and neurodegenerative diseases, leading to significant effort in developing strategies to limit its activation to intervene in these disorders. We previously showed that pharmacologic inhibition of the endoplasmic reticulum (ER)-localized protein disulfide isomerase PDIA1 suppresses NLRP3 activation and activity, identifying PDIA1 as a potential therapeutic target to mitigate hyperactive NLRP3 activity. Herein, we screen PDIA1 inhibitors to identify highly-potent compounds, including P1 and PACMA31, that pharmacologically target PDIA1 and block NLRP3 inflammasome assembly and activity. While sustained treatment with these PDIA1 inhibitors reduces THP1 viability, we show that acute treatment with these compounds is sufficient to both fully modify PDIA1 and inhibit NLRP3 inflammasome activity independent of any overt cellular toxicity. These results establish a treatment paradigm that can be exploited to develop highly-selective PDIA1 inhibitors to mitigate hyperactive NLRP3 inflammasome activity implicated in etiologically-diverse diseases.

## INTRODUCTION

The NLRP3 inflammasome is a multi-protein complex that functions to organize local and systemic immune signaling in response to diverse pathogen- and damage-associated signals.^1-4^ The activation of the NLRP3 inflammasome is generally associated with both priming and activation stimuli. Priming signals, such as pathogen associated molecular patterns (PAMPs) and damage-associated molecular patterns (DAMPs), activate the transcription factor NFκB to induce expression of inflammasome components such as NLRP3, ASC, and pro-caspase 1. Activation stimuli such as ATP or the pore forming toxin nigericin then induce assembly and posttranslational processing of these individual components into the active NLRP3 inflammasome.^1-4^ Once assembled and activated, the NLRP3 inflammasome promotes the caspase 1-dependent processing and release of pro-inflammatory cytokines including IL-1β and IL-18, which promote tissue inflammatory responses and neutralize pathogenic threats.^1-4^ The active NLRP3 inflammasome can also induce caspase 1-dependent cleavage of gasdermin D to promote pyroptotic cell death.^1-4^ While the NLRP3 inflammasome is an important component of the innate immune response, hyperactivity of this pathway can lead to tissue damage and cell death implicated in etiologically diverse diseases including cryopyrin-associated periodic syndrome (CAPS), diabetes, inflammatory disorders, and cardiac and cerebral ischemia/reperfusion.^5-14^ The involvement of hyperactive NLRP3 inflammasomes in these diseases has led to significant interest in developing strategies to restrict activity of this complex to mitigate the pathologic pro-inflammatory signaling associated with these disorders.^2,15,16^

Numerous small molecule-based strategies have been developed to inhibit NLRP3 inflammasome activation and activity. Compounds such as MCC950 and OLT1177 bind to the ATP binding pocket of NLRP3, preventing its activation and subsequent pro-inflammatory signaling.^17-19^ Further, NLRP3 is highly sensitive to electrophilic compounds, with covalent targeting of multiple different cysteine (Cys) residues across the NLRP3 structure inhibiting NLRP3 inflammasome assembly and activation.^20^ This has spurred considerable interest in developing electrophilic compounds such as itaconate and oridonin that selectively target NLRP3 to block inflammasome activity.^20-23^ Pharmacologic targeting of phosphoglycerate kinase 1 (PGK1) was also shown to inhibit NLRP3 inflammasome activation through a mechanism involving generation of the reactive metabolite methylglyoxal.^24^ Despite these efforts, no pharmacologic approaches are currently approved to inhibit pro-inflammatory NLRP3 activity in the context of human disease.

Recently, we showed that inhibition of the endoplasmic reticulum (ER) localized protein disulfide isomerase PDIA1 represents another strategy to inhibit NLRP3 activation.^25^ PDIA1 contains two redox-active Cys-X-X-Cys motifs, termed the a and a’ active sites, that function to promote and rearrange disulfides within proteins localized to the ER lumen.^26^ We demonstrated that pretreatment with metabolically activated compounds such as AA147 covalently targets PDIA1 in monocytes to inhibit NLRP3 inflammasome assembly and activity.^25^ Genetic depletion of *PDIA1* similarly reduced NLRP3 inflammasome activation and activity, confirming PDIA1 as an attractive therapeutic target for mitigating NLRP3 inflammasome activity.^25^ However, the requirement for pre-treatment with AA147 to induce PDIA1 modification to levels sufficient to inhibit NLRP3 inflammasome activity, combined with the low selectivity of this compound for PDIA1, has limited the continued development of AA147 for inhibiting NLRP3 inflammasomes through this mechanism.^25,27,28^

Many small molecules have been established as potent PDIA1 inhibitors. The widely used PDIA1 covalent inhibitor 16F16 was originally identified as a compound that could suppress PDIA1-dependent apoptosis induced by the toxic aggregation of proteins such as Aβ and huntingtin.^29^ The phenyl vinylsulfonate containing compound P1 was identified as a covalent inhibitor of human PDIA1 shown to reduce growth of multiple cancer cells and attenuate insulin aggregation.^30^ Further, PACMA31 was identified as an orally active covalent PDIA1 inhibitor that showed activity in cellular and in vivo models of ovarian cancer.^31^ Other compounds such as KSC34 have been developed as highly-selective PDIA1 inhibitors that selectively target the a active site of PDIA1 over the a’ active site.^32^ While these PDIA1 inhibitors have been shown to be beneficial in cellular and in vivo models of many different diseases, their therapeutic development has been largely limited by off-target activities associated with the chronic treatment of these electrophilic compounds.

Here, we screen commercially available covalent PDIA1 inhibitors and analogs of AA147 for their ability to inhibit NLRP3 inflammasome assembly and activity in THP1 monocytes. Through these efforts, we identified compounds P1 and PACMA1 as highly potent compounds that block activation of the pro-inflammatory NLRP3 inflammasome complex. However, we also found that chronic treatment with these compounds induced toxicity in LPS primed THP1 cells, likely through off target activities. To address this, we show that short, acute treatment with these compounds is sufficient to fully engage PDIA1 in these cells and suppress NLRP3 inflammasome assembly and activity. This demonstrates the potential for the continued development of covalent PDIA1 inhibitors with short tissue exposure times as a therapeutic strategy to mitigate pathologic NLRP3 hyperactivity implicated in etiologically diverse diseases.

## RESULTS & DISCUSSION

### Screening of small molecule PDIA1 inhibitors to suppress NLRP3 inflammasome assembly

We previously showed that pharmacologic PDIA1 inhibition using the metabolically activated compound AA147 blocks NLRP3 inflammasome assembly in THP1 monocytes.^25^ Here, we sought to identify other PDIA1 inhibitors with improved potency for inhibiting NLPR3 inflammasome assembly in these cells. Towards that aim, we screened a panel of AA147 analogs^28,33^ and established PDIA1 inhibitors in dose response for their ability to block NLRP3 assembly in THP1 ASC-GFP cells. We pretreated these cells with the priming signal LPS for 16 h and then added compound for an additional 16 h (**Fig. 1A**). We then then induced NLRP3 inflammasome assembly by treatment with the activating stimulus nigericin. NLRP3 inflammasomes can then be followed by the formation of ASC-GFP specks 2 h after nigericin treatment quantified by high content imaging (**Fig. S1A**).^20,24,25^ Our screen identified 3 AA147 analogs (AA55, AA62, and AA66) and 3 commercially available PDIA1 inhibitors (P1, PACMA31, and 16F16) as the most potent inhibitors of ASC-GFP speck formation (**Fig. 1A, Fig. S1B**). We rescreened these compounds for their dose-dependent inhibition of ASC-GFP speck formation in THP1 ASC-GFP cells. This showed that 16 h treatment with AA62 and AA66 had IC_50_s for ASC-GFP speck formation of 4.94 µM or 6.17 µM, respectively, while AA55 did not reproducibly reduce ASC-GFP speck formation (**Fig. S1C-E**). However, 6 h pretreatments with 16F16 (IC_50_ = 2.83 µM), P1 (IC_50_ = 890 nM), and PACMA31 (IC_50_ = 146 nM) all reduced ASC-GFP speck formation with higher potency than our AA147 analogs (**Fig. 1B-D**). Thus, we selected P1 and PACMA31 as the top two compounds from this screen for further study.

**Figure 1.**
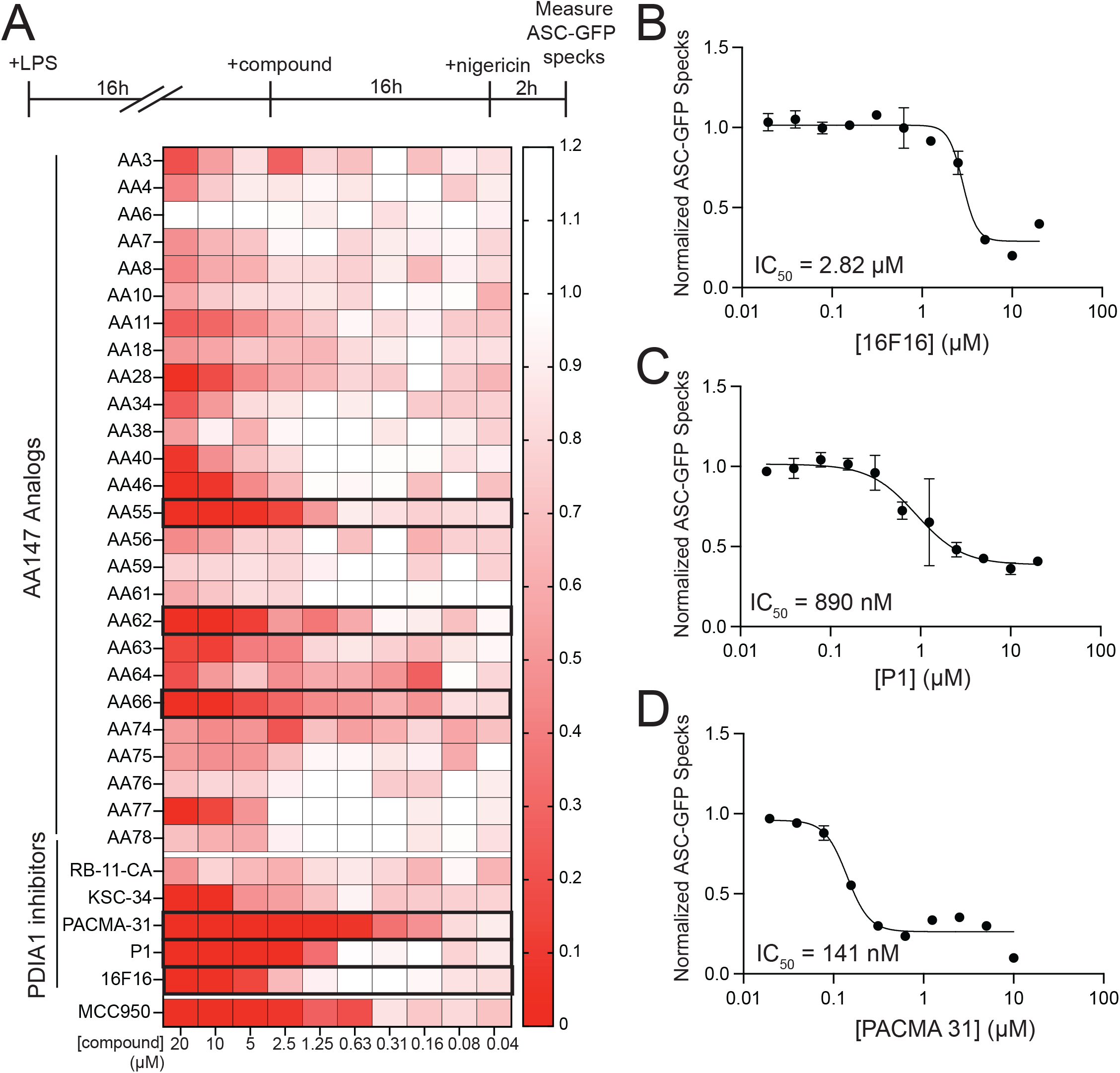
Identification of PDIA1 inhibitors with improved potency for inhibiting ASC-GFP speck formation. **A.** Heat map showing normalized ASC-GFP speck formation quantified from microscopy images of THP1 ASC-GFP cells that were pre-treated for 16 h with LPS (1 µg/mL) and the indicated compound (10 µM) for 16 h and then stimulated with nigericin (10 µM) for 2 h. The NLRP3 inhibitor MCC950 is shown as a control. **B-D**. Normalized ASC-GFP speck formation quantified from microscopy images of THP1 pretreated for 16 h with LPS (1 µg/mL) and stimulated for 2 h with nigericin (10 µM). Compounds were added at the indicated concentration 6 h prior to nigericin treatment. Error bars show SEM for n=3 replicates. The IC_50_ for inhibition of ASC-GFP speck formation is shown.

### P1 and PACMA31 rapidly engage PDIA1 to inhibit NLRP3 inflammasome assembly

P1 contains an electrophilic vinyl sulfonate that enables covalent modification of the active site Cys residues on PDIA1, as well as an alkyne moiety suitable for monitoring PDIA1 engagement by P1 using click chemistry (**Fig. S1B**).^30^ We treated LPS-primed THP1 monocytes with P1 for 6 h and monitored covalent protein engagement by appending a rhodamine azide to P1-modified proteins using click chemistry. This identified a prominent band at ∼57 kDa, which corresponds to the molecular weight of PDIA1 (**Fig. 2A, Fig. S2A**). Using an analogous experimental approach where we appended a biotin-azide to P1 modified proteins to allow streptavidin-based affinity isolation, we showed that P1 dose-dependently engages PDIA1 (**Fig. 2B**). We also observed engagement of other ER-localized PDIs including PDIA4 and PDIA6, which are often observed with PDIA1 inhibitors owing to the similarities in the active sites across these enzymes. Importantly, we do not observe modification of NLRP3 in these assays, indicating that these compounds do not inhibit NLRP3 inflammasome activation through direct covalent targeting of NLRP3. While PACMA31 does have an alkyne handle, this moiety is lost upon covalent protein modification, precluding our use of click chemistry to directly monitor protein engagement by this compound.^31^ However, we were able to demonstrate that co-treatment with PACMA31 blocked P1-dependent labeling of PDIA1 and other labeled proteins (**Fig. 2A,B**). These results confirm that both P1 and PACMA31 covalently engage PDIA1 in LPS primed THP1 cells.

**Figure 2.**
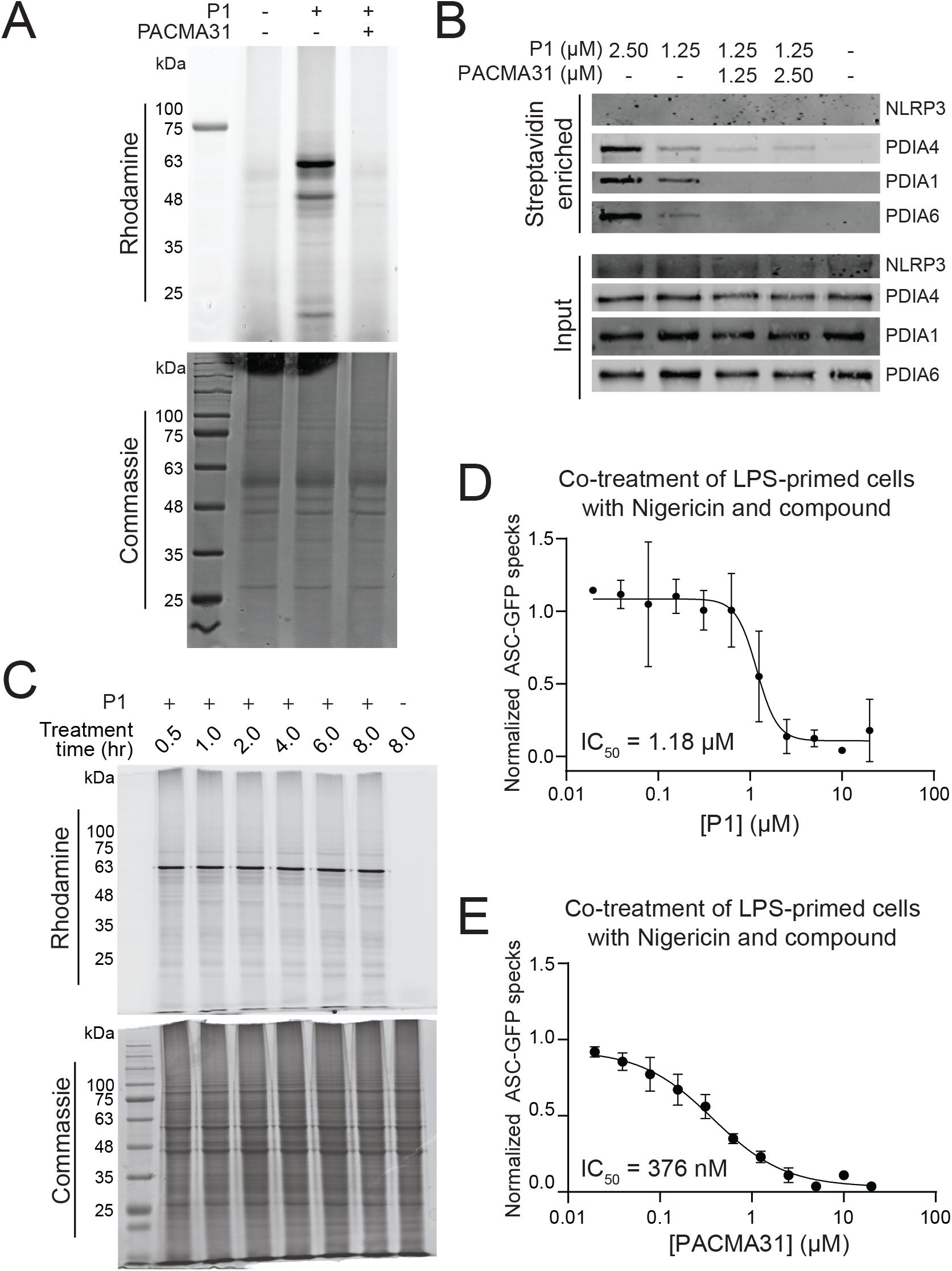
P1 and PACMA31 rapidly modify PDIA1 in THP1 cells. **A.** Fluorescent and Coomassie stained SDS-PAGE gels of lysates prepared from THP1 null cells primed for 16 h with LPS (1 µg/mL) treated with vehicle, P1 (500 nM), and/or PACMA31 (20 µM) for 6 h. P1 modified proteins were identified by click-dependent modification of the alkyne with rhodamine-azide. **B.** Immunoblot of input and streptavidin-affinity purified P1 modified proteins in THP1 null cells primed for 16 h with LPS (1 µg/mL) treated with the indicated concentration of P1 and/or PACMA31 for 6 h. P1 modified proteins were modified with biotin through click-dependent addition of a biotin-azide to the alkyne moiety of P1. **C**. Fluorescent and Coomassie stained SDS-PAGE gels of lysates prepared from THP1 null cells primed for 16 h with LPS (1 µg/mL) treated with P1 (2.5 µM) for the indicated time. P1 modified proteins were identified by click-dependent modification of the alkyne with rhodamine-azide. **D,E**. Normalized ASC-GFP speck formation quantified from fluorescent images of THP1 ASC-GFP cells that were pretreated with LPS (1 µg/mL) and then co-treated with nigericin (10 µM) and the indicated concentration of P1 (**D**) or PACMA31 (**E**) for 2 h. Error bars show SEM for n=3 replicates. The IC_50_ for inhibition of ASC-GFP speck formation is shown.

Next, we sought to define the rate of PDIA1 engagement by covalent inhibitors such as P1. We treated LPS primed THP1 cells with P1 for varying lengths of time and then monitored covalent engagement by appending a rhodamine azide to modified proteins by click chemistry. Intriguingly, treatment with P1 for as little as 0.5 h was sufficient to fully engage proteins in these cells, with no significant increase observed with longer treatment (**Fig. 2C**). This suggested that inhibition of NLRP3 inflammasome assembly in these cells could be observed with compound pre-treatments of <0.5 h. Consistent with this, pretreatment of LPS-primed THP1 ASC-GFP monocytes with P1 or PACMA31 for 2 or 4 h was sufficient to block ASC-GFP speck formation induced by nigericin treatment (**Fig. S2B-E**). Moreover, we found that co-treatment of LPS-primed THP1 monocytes with nigericin and either P1 or PACMA31 potently inhibited ASC-GFP speck formation in THP1 ASC-GFP monocytes (**Fig. 2D,E**). These results show that PDIA1 is rapidly engaged by P1 and PACMA31, allowing for efficient inhibition of NLRP3 activation independent of a pretreatment.

### Chronic treatment with P1 or PACMA31 is toxic to THP1 monocytes

We next sought to determine the potential for P1 and PACMA31 to inhibit downstream aspects of NLRP3 inflammasome activation such as IL1β secretion and pyroptotic cell death. We initially monitored IL1β secretion from THP1 monocytes pretreated with LPS and then co-treated with the activating stimulus ATP and either P1 or PACMA31 for 24 h. We collected media from these cells and added this media to HEK-Blue-IL1β cells – a cell line that stably expresses the IL1β responsive TLR receptor and the NFκB/AP1-inducible secreted alkaline phosphatase (SEAP) reporter.^20,24,25^ Thus, mature IL1β secreted from THP1 cells into conditioned media can be monitored by the secretion of SEAP from HEK-Blue-IL1β cells. We found that treatment with P1 or PACMA31 dose-dependently reduced IL1β secretion from THP1 cells, as measured by this HEK-Blue-IL1β SEAP secretion assay (**Fig. 3A**). Next, we monitored pyroptotic cell death in THP1 monocytes pretreated for 16 h with LPS and then 2 h with P1 or PACMA31. We then added nigericin for 2 h prior to monitoring cellular viability with CellTiter-Glo. Cellular viability was monitored by CellTiter-Glo. Intriguingly, both P1 and PACMA31 rescued viability of LPS-primed and nigericin treated THP1 cells at concentrations near the IC_50_ for inflammasome inhibition (P1 ∼1 µM or PACMA31 ∼300 nM; **Fig. 3B**), indicating that these compounds reduced pyroptosis. However, at higher concentrations, we observed significant reductions in viability under these conditions. This suggests that P1 and PACMA1 show toxicity in THP1 cells at elevated concentrations. Consistent with this, treating LPS-primed THP1 cells with P1 or PACMA31 for 16 h dose-dependently reduced THP1 viability, as measured by CellTiter-Glo (**Fig. 3C**). However, this toxicity was abrogated in cells treated with these compounds for only 4 h (**Fig. S3**). These results indicate that chronic treatment with P1 or PACMA31 can induce toxicity in THP1 monocytes.

**Figure 3.**
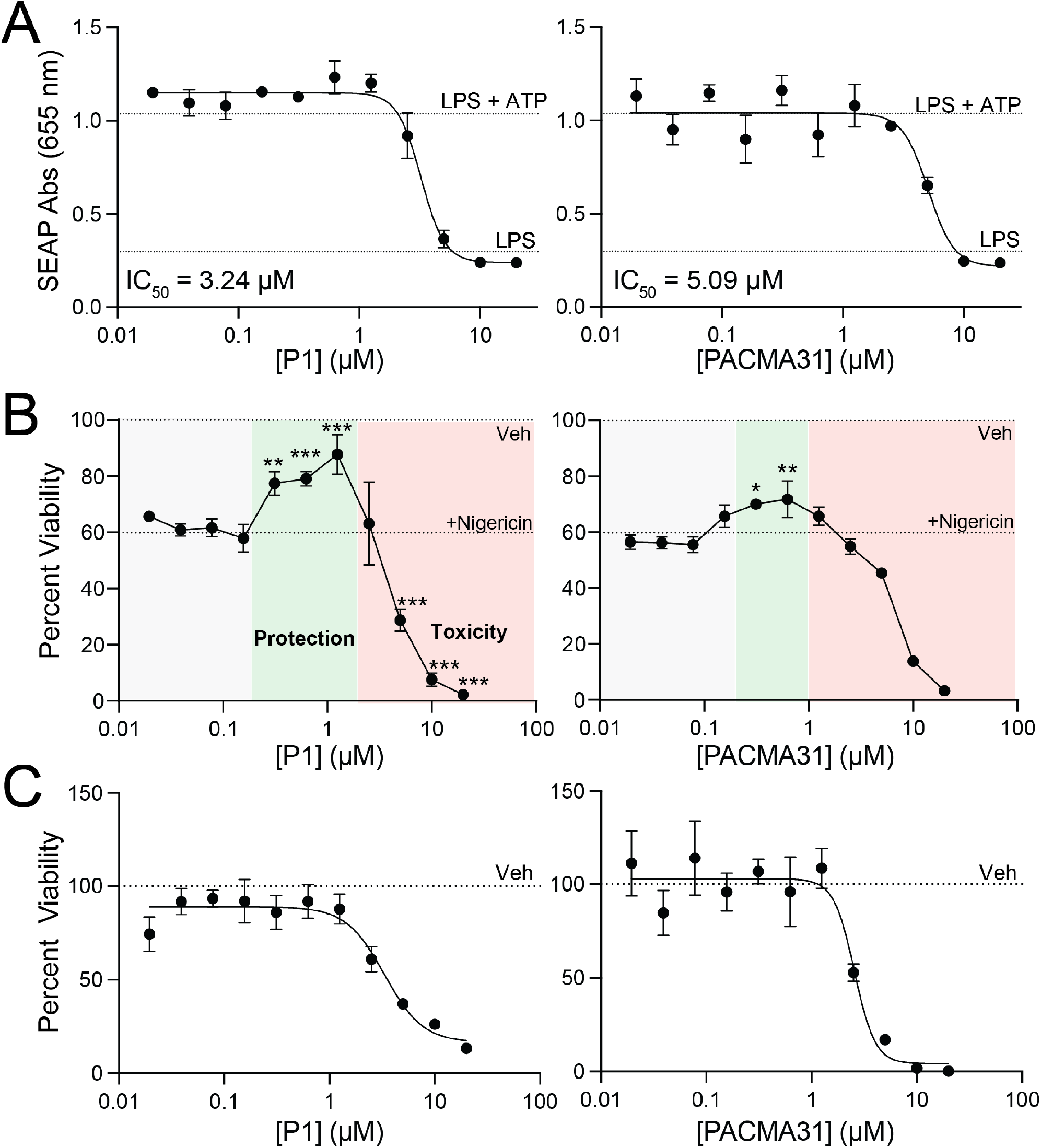
Chronic treatment of THP1 cells with P1 and PACMA31 reduces cell viability. **A.** Secreted IL1β, measured by HEK-Blue-IL1β, from THP1 null cells primed for 16 h with LPS (1 µg/mL) and then co-treated for 24 h with the indicated concentration of P1 (left) or PACMA31 (right) and ATP (5 mM). Error bars show SEM for n=3 replicates. The IC_50_ for inhibition of IL1β secretion is shown. **B.** Viability of THP1 null cells primed for 16 h with LPS (1 µg/mL) and treated with nigericin (10 µM) for 2 h. The indicated concentration of P1 (left) or PACMA31 (right) were added 2 h prior to nigericin treatment. Error bars show SEM for n=3 replicates. *p<0.05, **p<0.01, ***p<0.005 for one-way ANOVA relative to cells treated with nigericin alone. **C**. Viability of THP1 null cells pretreated for 16 h with LPS (1 µg/mL) and then treated for an additional 16 h with the indicated concentration of P1 (left) or PACMA31 (right). Error bars show SEM for n=3 replicates.

### Acute exposure to P1 or PACMA31 is sufficient for effective PDI engagement and NLRP3 inhibition independent of toxicity

Acute treatments of THP1 cells with P1 for <0.5 h is sufficient for effective engagement of protein targets such as PDIA1 (**Fig. 2C**). Further, we found that P1 modified proteins in THP1 monocytes are not significantly reduced following a 6 h treatment with cycloheximide (CHX), indicating that these modified proteins are relatively stable (**Fig. S4A**). These results suggest that acute exposure to P1 or PACMA31 should be sufficient to fully engage PDIA1 and maintain NLRP3 inflammasome inhibition, providing an opportunity to prevent the toxicity observed upon longer incubations. To test this, we treated LPS primed THP1 monocytes with P1 or PACMA31 for 1 h and then replaced the media with fresh media lacking compound for 24 h and monitored viability by CellTiter-Glo (**Fig. 4A**). As a control, we also prepared cells where P1 or PACMA31 was re-added to the replacement media to mimic the chronic treatment with these compounds shown to induce toxicity in these THP1 monocytes (see **Fig. 3C**). We found that cells exposed to compound for only 1 h demonstrated viability at levels comparable to untreated, LPS stimulated THP1 monocytes and greater than that observed for cells exposed to compound for the entire 24 h period (**Fig. 4B**). Next, we wanted to determine if this acute treatment would also be sufficient to inhibit NLRP3 inflammasome assembly and activity. To test this, we treated LPS-stimulated THP1 ASC-GFP monocytes with P1 or PACMA31 for 1 h, then replaced the media with fresh media with or without compound for an additional 1 h. We then stimulated these cells with nigericin and monitored ASC-GFP speck formation. We found that the 1 h treatment was sufficient to effectively inhibit ASC-GFP speck formation even after incubation in compound-free media for an additional 1 h (**Fig. 4C**). Importantly, this inhibition was still maintained when the incubation time in compound-free media following 1 h of treatment was extended to 8 h (**Fig. 4D**). However, this effect was lost as the incubation period was increased to 16 h (**Fig. S4B**). Apart from ASC-GFP speck formation, we also found that the 1 h treatment with compound was sufficient to reduce IL1β secretion from THP1 cells treated with LPS and ATP, as measured by SEAP secretion from HEK-Blue-IL1β cells (**Fig. 4E**). Further, we probed the potential for acute treatment with P1 or PACMA31 to decrease IL1β secretion from human peripheral blood mononuclear cells (PBMCs), where we found treatment with LPS alone, independent of activating stimulus, was sufficient to increase IL1β secretion (**Fig. 4F**). Interestingly, acute treatment with either P1 or PACMA1 decreased IL1β secretion from PBMCs. These results demonstrate that acute treatments with pharmacologic PDIA1 inhibitors such as P1 or PACMA31 is sufficient to inhibit NLRP3 inflammasome assembly and activity independent of toxicity associated with longer treatment with these covalent compounds.

**Figure 4.**
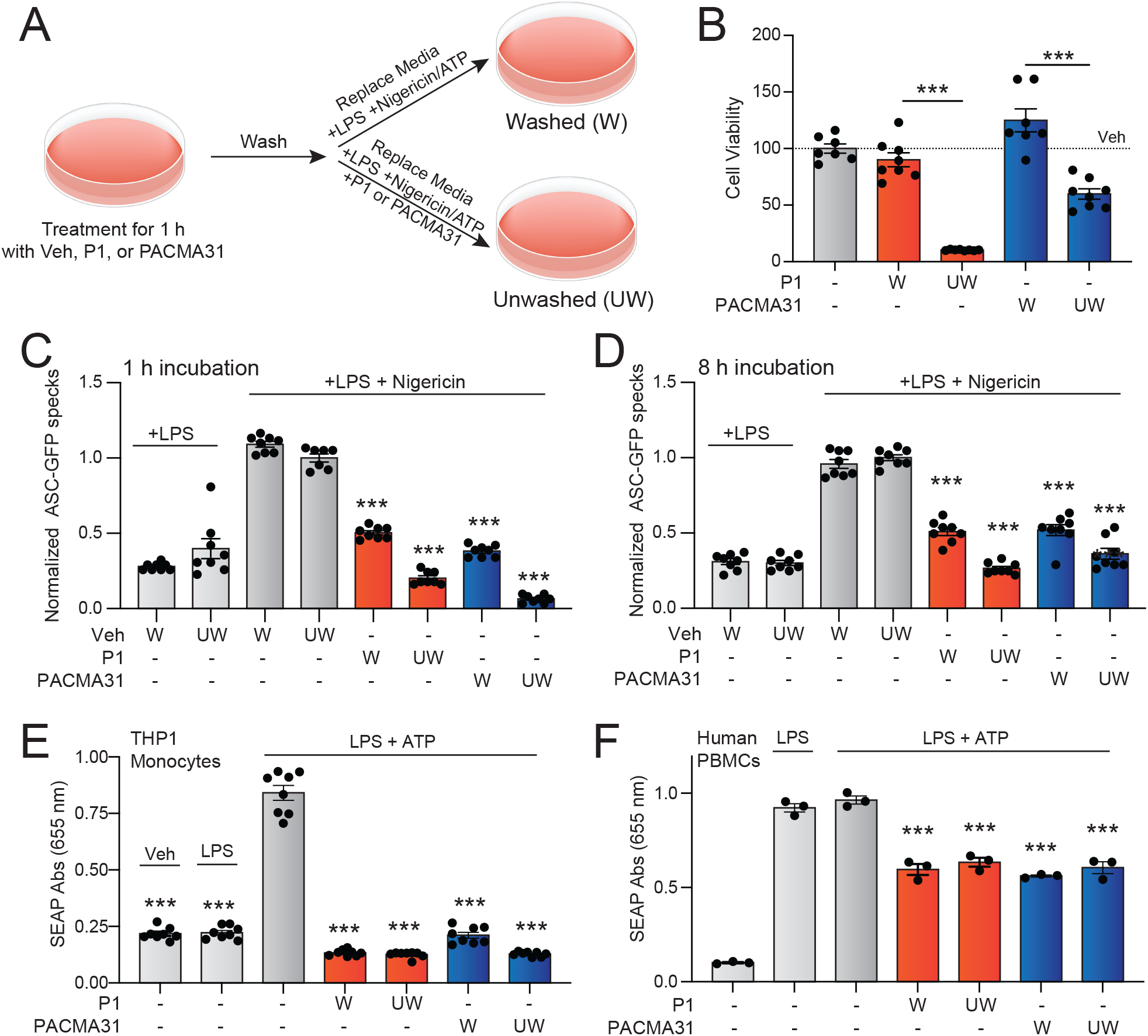
Acute treatment with P1 or PACMA31 blocks NLRP3 inflammasome activation independent of cellular toxicity. **A.** Protocol for acute treatment with P1 or PACMA31. Briefly, THP1 cells are treated for 1 h with P1 or PACMA31. The cells are then washed, and the media is replaced with fresh media both without (‘washed’) or with (‘unwashed’) P1 or PACMA1 added. **B**. Viability of THP1 null cells primed for 16 h with LPS (1 µg/mL) and then treated for 1 h with P1 (10 µM) or PACMA31 (10 µM). The cells were then washed, and the media was replaced with LPS in the absence (washed, W) or presence (unwashed, UW) of P1 or PACMA31. Viability was measured after 24 h. Error bars show n=8 replicates. **C,D.** ASC-GFP speck formation from THP1 ASC-GFP pretreated for 16 h with LPS (1 µg/mL) and treated for 1 h with P1 (5 µM) or PACMA31 (5 µM). The cells were then washed, and the media was replaced with LPS in the absence (washed, W) or presence (unwashed, UW) of P1 or PACMA31. Nigericin (10 µM) was added for 2 h prior to imaging of ASC specks at the end of 3 h (**C**) or 8 h (**D**) compound incubation. Error bars show SEM for n=8. **E**. Secreted IL1β, measured by HEK-Blue-IL1β, from THP1 null cells primed for 16 h with LPS (10 µM) and treated for 1 h with P1 (10 µM) or PACMA31 (10 µM). The cells were then washed, and the media was replaced with LPS in the absence (washed, W) or presence (unwashed, UW) of P1 or PACMA31. After 1 h, ATP (5 mM) was added for 24 h prior to media collection and analysis of secreted IL1β by HEK-Blue-IL1β. Error bars show SEM for n=8. **F.** Secreted IL1β, measured by HEK-Blue-IL1β, from human PBMCs primed for 16 h with LPS (10 µM) and treated for 1 h with P1 (10 µM) or PACMA31 (10 µM). The cells were then washed, and the media was replaced with LPS in the absence (washed, W) or presence (unwashed, UW) of P1 or PACMA31. After 1 h, ATP (5 mM) was added for 24 h prior to media collection and analysis of secreted IL1β by HEK-Blue-IL1β. Error bars show SEM for n=8. ***p<0.005 for two-way ANOVA relative to vehicle treated cells (**B**) or vehicle-treated cells administered LPS and nigericin/ATP (**C-F**).

## CONCLUDING REMARKS

Herein, we identify P1 and PACMA31 as highly potent PDIA1 inhibitors that block NLRP3 inflammasome assembly and activity. Further, we demonstrate that acute treatments with these compounds are sufficient to sustain NLRP3 inflammasome inhibition independent of the toxicity associated with chronic treatment with these compounds. These results demonstrate the potential for PDIA1 targeting to mitigate pathologic NLRP3 hyperactivity implicated in etiologically diverse diseases. Moreover, our results establish a therapeutic paradigm that can help further develop PDIA1 inhibitors for this purpose. We show that acute treatment with PDIA1 inhibitors such as P1 and PACMA31 is sufficient to effectively and stably engage PDIA1 to levels suitable for inhibiting NLRP3 inflammasome activity. Since our results indicate that longer treatments lead to cell toxicity, without increasing PDIA1 labeling, it is likely that this toxicity results from off-target activities of these electrophilic compounds that accumulate over time. Accordingly, compounds with good bioavailability capable of rapidly and selectively engaging PDIA1 that are rapidly cleared from systemic circulation may offer unique opportunities to target PDIA1 to inhibit NLRP3 inflammasome activity independent of dose-limiting toxicity. Thus, as new PDIA1 inhibitors are developed, optimization of the pharmacokinetic and pharmacodynamic properties of these compounds will likely allow for improved safety and activity for this purpose. Ultimately, our work has highlighted the therapeutic potential of pharmacologically targeting PDIA1 for ameliorating NLRP3 hyperactivity implicated in diverse diseases, spurring the continued development of highly selective compounds with the PK/PD properties suitable for continued translational development.

## MATERIALS AND METHODS

### Cell culture

THP1 null cells (Invivogen; generous gift from John Griffin, TSRI) and THP1 ASC-GFP cells (Invivogen; generous gift from Calibr, TSRI) were maintained in RPMI 1640 (Lifetech cat. 11875135) supplemented with 10% heat-inactivated fetal bovine serum (FBS) (Lifetech cat. 10082147), 100 μg/ml Normocin (Invivogen cat. ant-nr-1), and Pen-Strep (100 μg/ml). HEK-Blue-IL1β reporter cells (Invivogen; cat # hkb-il1bv2) were maintained in DMEM supplemented with 10% heat-inactivated FBS (Lifetech cat. 10082147), 2 mM glutamine, and Pen-Strep (100 μg/ml). Peripheral Blood Mononuclear (PBMCs) was isolated from whole blood collected by Scripps Normal Blood Donor Service (NBDS). Briefly, whole blood was collected and processed for plasma and peripheral blood mononuclear cells (PBMCs) using Ficoll-Plaque Plus (GE Healthcare, Chicago, IL) and Leucosep (Greiner Bio-One, Kremsmünster, Austria) tubes, following the manufacturers’ instructions. PBMCs were maintained in RPMI 1640 Medium with GlutaMAX™ Supplement (Lifetech cat. 61870036), 10% FBS, Pen-Strep (100 μg/ml), and 10 mM HEPES.

### Compounds and Treatments

Ultrapure LPS from E. coli O111:B4 (Invivogen cat # tlrl-3pelps) at 1 µg/mL was used to induce the NFκB transcription and NLRP3 inflammasome priming in all experiments. Inflammasome activation and assembly were induced with a treatment of 5 mM ATP (Cytiva cat. 45001323) in water or 10 μM nigericin (Invivogen cat. tlrl-nig) in 100% 200-proof Ethanol. The AA147 analogs were previously reported.^28,33^ 16F16 was purchased from Sigma-Aldrich (cat. SML0021). KSC-34 was purchased from Probechem (cat. PC-63459). RB-11-CA was a kind gift from the Weerapana Lab at Boston College. P1 was purchased from WuXi AppTec. PACMA31 was purchased from Med Chem Express (cat. HY-100433). Cycloheximide was purchased from Fisher (cat. AAJ66901-03). Compounds were suspended in dimethyl sulfoxide (DMSO). MCC950 was purchased from Selleck Chem (cat. S7809) and was resuspended in water.

### Antibodies

The primary antibodies were diluted as noted below in 5% bovine serum albumin (BSA) and 0.1% Sodium Ascorbate in TBS and incubated overnight: NLRP3 (1:1000; Cell Signaling cat. 15101), PDIA1 (1:1000, Gene Tex cat. GTX101468), PDIA4/Erp72 (1:1000, Protein Tech cat. 14712-1-AP), and PDIA6 (1:1000, Protein Tech cat. 66669-1-Ig).

### ASC-GFP specks assay

THP1 ASC-GFP cells were primed with LPS for 16 h. Cells were plated at the density of 20,000 cells/well in a 384 well plate and treated with compounds at indicated doses and time. Next, cells were stimulated with nigericin (10 µM) and stained with Hoechst 33342 (Thermo cat. H3570) for 2h. Cellomics Cell Insight imaging reader (Thermo) with a 10x air lens was used to capture one image of GFP and nuclei per well. Images were analyzed for ASC-GFP speck level of each condition, as before.^20,24,25^ The total amount of cells was calculated from the number of Hoechst-stained nuclei. The specks were detected and calculated using an algorithm within the Cell Insight software based on both signal intensity and area size. Numerical results from the analyzed images were later exported for analysis. Cells treated with vehicle and nigericin were used as maximum specks formation control. Cells treated with vehicles but not treated with nigericin were used as minimum specks formation control. Relative ASC-GFP specks formation was then calculated based on the maximum and minimum specks formation controls.

### Pyroptotic cell death assay

THP1 nulls cells were primed with LPS, pretreated with covalent PDI inhibitors, and activated with nigericin as previously described in the ASC-GFP Speck Assay. CellTiter-Glo (CTG) (Promega cat. PRG7572; diluted 1:6) was added to the stimulated cells at 1:3 the volume of growth medium. Luminescence was measured via Tecan Infinite M200 Pro Microplate Reader. Cells treated with vehicle and nigericin were used as positive control (maximum cell death). Cells treated with vehicle but not treated with nigericin were used as negative control (maximum viability and minimal cell death). Percent viability at each concentration was calculated relative to the luminescence of negative control.

### HEK-Blue-IL1β secretion assay

THP1 nulls cells were primed with LPS for 16 h. Cells were plated at the density of 40,000 cells/well in a 384 well plate and treated with compounds at indicated doses and time. Next, cells were stimulated with ATP (5 mM) for 24 h. Low speed centrifugation was used to collect the conditioned media in the supernatant. Each conditioned medium was added directly to the HEK-Blue IL-1β Reporter Cells at 10,000 cells/well in a 384 well plate for 16 h. Quanti-blue (Invivogen cat. rep-qbs) was used to detect the levels of the secreted alkaline phosphatase reporter in the HEK-Blue-IL1β cells. Absorbance of SEAP at 655 nm was measured using SPECTRAmax PLUS 384 (Molecular Devices) plate reader.

### Cell viability assay

THP1 nulls cells were primed with LPS, pretreated with covalent PDI inhibitors, and incubated until indicated time. CTG was used to measure cell death caused by the compounds at each treatment time as previously described in the pyroptotic cell death assay. Cells treated with vehicle were used as negative control (minimum cell death, 100% viability) and were used to calculate percent viability for other conditions.

### Protocol for washing THP1 monocytes to remove PDI inhibitors

LPS-primed THP1 ASC-GFP cells, THP1 nulls cells, or PBMCs were treated with P1 or PACMA31 for 1 h. The compounds were then removed from the cells by 3 cycles of centrifugation and growth medium replacement. LPS was reapplied if the cells were previously primed with LPS. Unwashed controls were created by treating the wash cells with the same dose of P1 or PACMA31. Both the washed and the unwashed cells were then seeded for further analyses with ASC-GFP specks assay, cell viability assay, and IL1b secretion assay.

### Rhodamine labeling by Click Chemistry

Cell lysates were prepared from various treatments of LPS-primed THP1 cells by sonication in RIPA buffer (50 mM Tris, pH 7.5, 150 µM NaCl, 1% Triton X-100, 0.5% sodium deoxycholate, 0.1% SDS, and protease inhibitor cocktail (Pierce cat. A32955)). Rhodamine labeling was performed in a 200 µL reaction containing 300 µg of total protein lysate. Each reaction contained a final concentration of 0.8 mM CuSO4, 1.6 mM BTTAA (Sigma cat. 906328-100MG), 5mM Sodium Ascorbate (Fisher cat. A4034), and 0.1 mM TAMRA azide (Santa Cruz cat. sc-482005). Reactions were incubated at 37°C for 2 h. Labeled proteins were purified by MeOH precipitation and 4:1:1 MeOH: CHCl_3_: water washes. Proteins were resuspended in 3 M Urea in DPBS with 1% SDS. Samples were denatured by boiling in 1× Laemmli buffer with 100 mM dithiothreitol (DTT) before being separated by SDS-PAGE. Gel was then stained with 0.1% Coomassie Blue R250 in 10% acetic acid and 50% methanol. Both rhodamine fluorescence and Coomassie staining were imaged on a ChemiDoc Imaging Systems (BioRad).

### Biotin labeling by Click Chemistry and Affinity Purification

Cell lysates were prepared from various LPS-primed THP1 cells by sonication in RIPA buffer. Biotin labeling was performed in a 200 µL reaction containing 300 µg of total protein lysate from LPS-primed THP1 cells treated with compounds. Biotin labeling was performed as previously described in a 200 µL reaction containing 300 µg of total protein lysate.^20^ Each reaction contained a final concentration of 0.8 mM CuSO4, 1.6 mM BTTAA (Sigma cat. 906328-100MG), 5mM Sodium Ascorbate (Fisher cat. A4034), and 0.1 mM Biotin-azide (Med Chem Express cat. HY-129832). Reactions were incubated at 37°C for 2 h. Labeled proteins were purified by MeOH precipitation and 4:1 MeOH: CHCl_3_ washes. Proteins were resuspended in 0.75 M Urea in DPBS with 0.25% SDS. Streptavidin agarose slurry (Thermo cat. 20349) were added and incubated overnight at 4°C to enrich biotinylated proteins. Elution of biotin-labeled proteins were performed by washing the beads with 0.1% SDS in DPBS and DPBS then boiling the beads with 1x Laemmli buffer + 100 mM DTT. Immunoprecipitation inputs were also denatured with 1x Laemmli buffer + 100 mM DTT and boiled before being separated alongside the enriched samples by SDS-PAGE. Separated samples were transferred onto nitrocellulose membranes (Bio-Rad cat. 1620112). Membranes were then incubated overnight at 4 °C with primary antibodies with dilutions as noted. Membranes were washed in TBS-T, incubated with the species-appropriate IR-Dye conjugated secondary antibodies diluted at 1:10,000, and analyzed with the Odyssey Infrared Imaging System (LI-COR Biosciences).

## AUTHOR CONTRIBUTIONS

C.B., C.S., designed research; C.B., C.S., P.B., and P.C. performed research and analyzed data; C.B., M.J.B., and R.L.W. wrote the paper. M.J.B., and R.L.W. received funding and oversaw the project.

## ACKNOWLEDGEMENTS

We thank Mansun Law (Scripps) for experimental advice related to this project and Jessica Rosarda (USUHS) for the critical reading of this manuscript. This work was supported by the National Institutes of Health (AG046495 and DK107604 to RLW) and the Royal Thai Government Fellowship (to CB).

---

**Figure S1.**
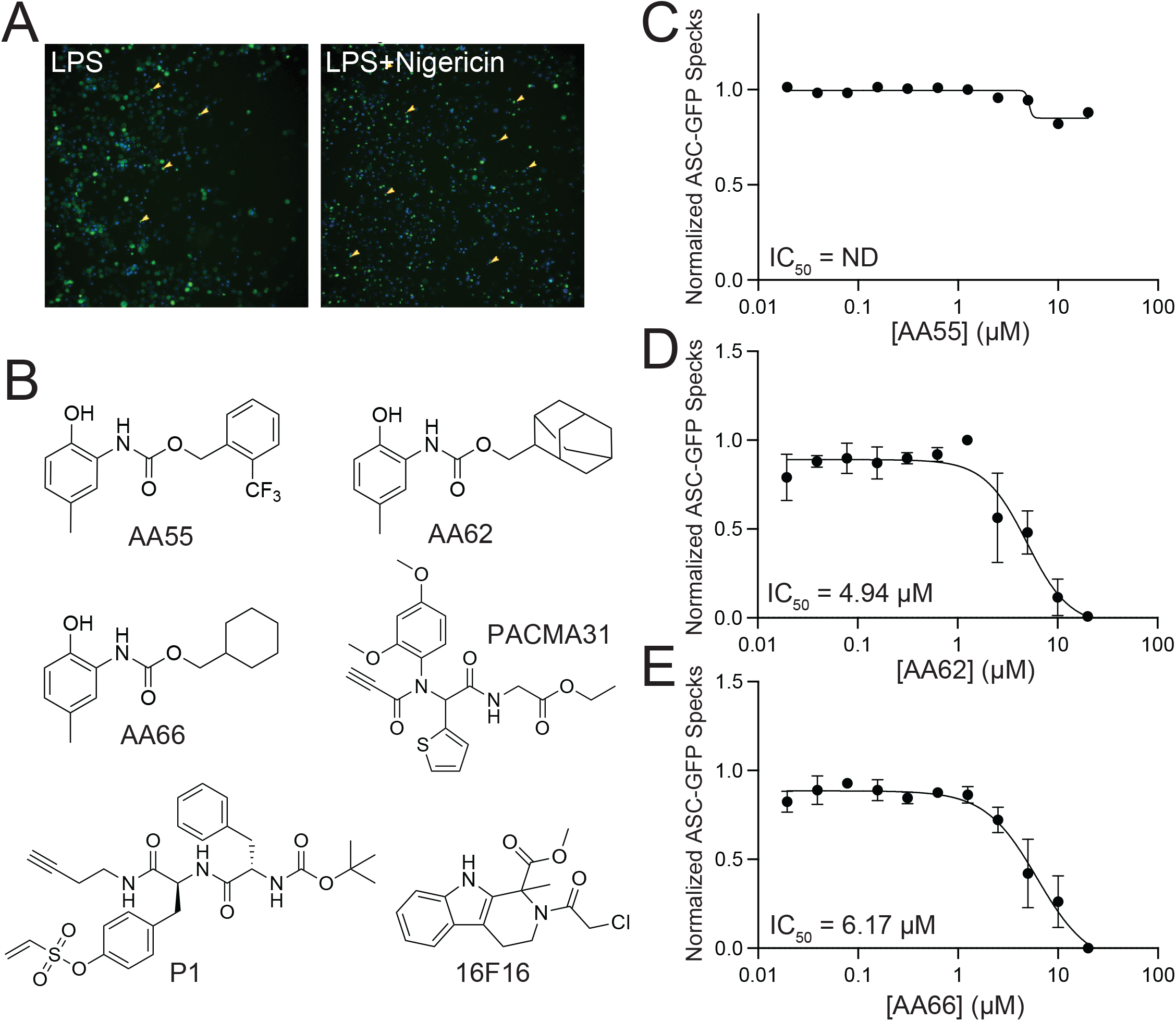
(Supplement to Figure 1) Identification of PDIA1 inhibitors with improved potency for inhibiting ASC-GFP speck formation. **A.** Representative fluorescent images of THP1 ASC-GFP cells pretreated with LPS (1 µg/mL) for 16 h and then treated with nigericin (10 µM) for 2 h. Arrows identify ASC-GFP specks. **B**. Structures of prioritized AA147 analogs and commercial PDIA1 inhibitors identified in the screen showed in **Fig. 1A. C-E.** Normalized ASC-GFP speck formation quantified from fluorescent images of THP1 ASC-GFP cells that were pre-treated for 16 h with LPS (1 µg/mL) and the indicated compound (10 µM) for an additional 16 h. We then stimulated with nigericin (10 µM) for 2 h prior to monitoring ASC-GFP specks. Error bars show SEM for n=3 replicates. The IC_50_ for inhibition of ASC-GFP speck formation is shown.

**Figure S2.**
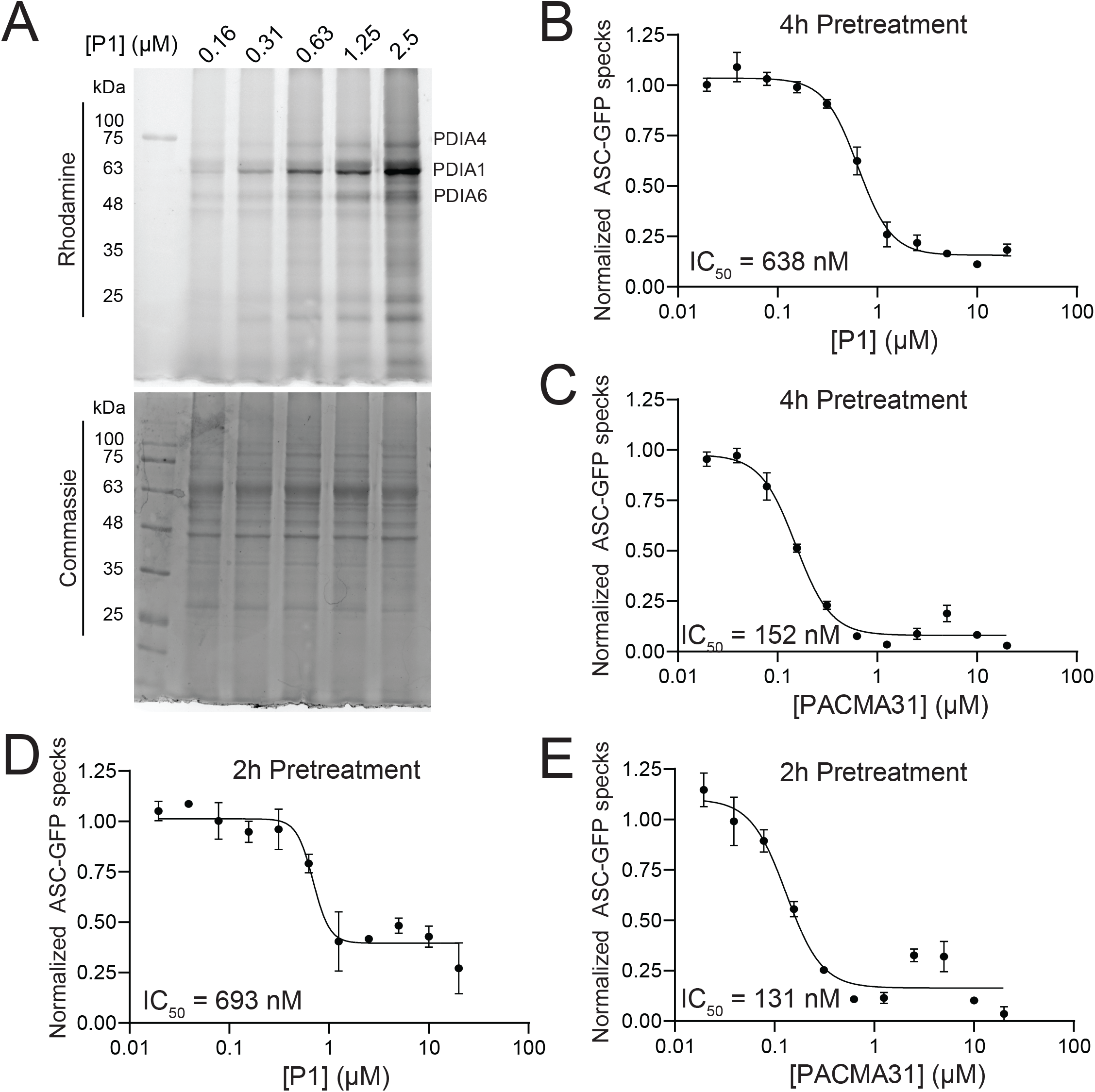
(Supplement to Figure 2). P1 and PACMA31 rapidly modify PDIA1 in THP1 cells. **A.** Fluorescent and Coomassie stained SDS-PAGE gels of lysates prepared from THP1 null cells primed for 16 h with LPS (1 µg/mL) treated with the indicated concentration of P1 for 6 h. P1 modified proteins were identified by click-dependent modification of the alkyne with rhodamine-azide. **B,C**. Normalized ASC-GFP speck formation quantified from microscopy images of THP1 pretreated for 16 h with LPS (1 µg/mL) and stimulated for 2 h with nigericin (10 µM). Compounds were added at the indicated concentration 4 h prior to nigericin treatment. Error bars show SEM for n=3 replicates. The IC_50_ for inhibition of ASC-GFP speck formation is shown. **D,E**. Normalized ASC-GFP speck formation quantified from microscopy images of THP1 pretreated for 16 h with LPS (1 µg/mL) and stimulated for 2 h with nigericin (10 µM). Compounds were added at the indicated concentration 2 h prior to nigericin treatment. Error bars show SEM for n=3 replicates. The IC_50_ for inhibition of ASC-GFP speck formation is shown.

**Figure S3.**
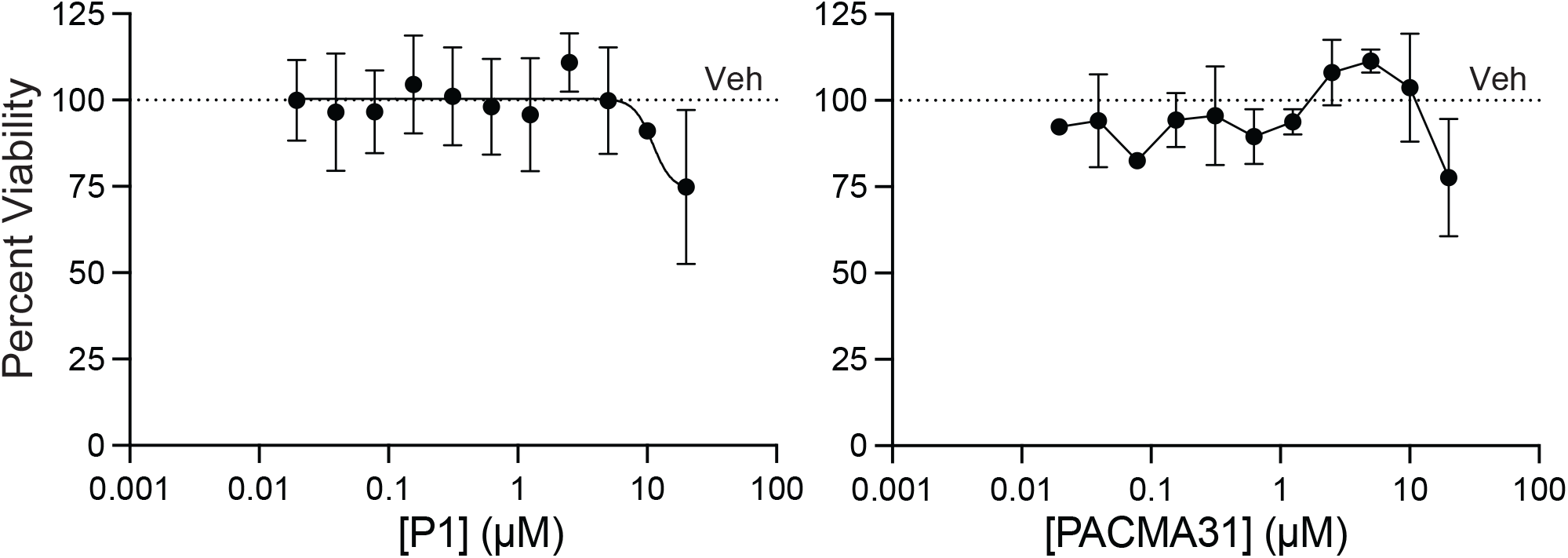
(Supplement to Figure 3). Chronic treatment of THP1 cells with P1 and PACMA31 reduces cell viability. Viability of THP1 null cells pretreated for 16 h with LPS (1 µg/mL) and then treated for an additional 4 h with the indicated concentration of P1 (left) or PACMA31 (right). Error bars show SEM for n=3 replicates.

**Figure S4.**
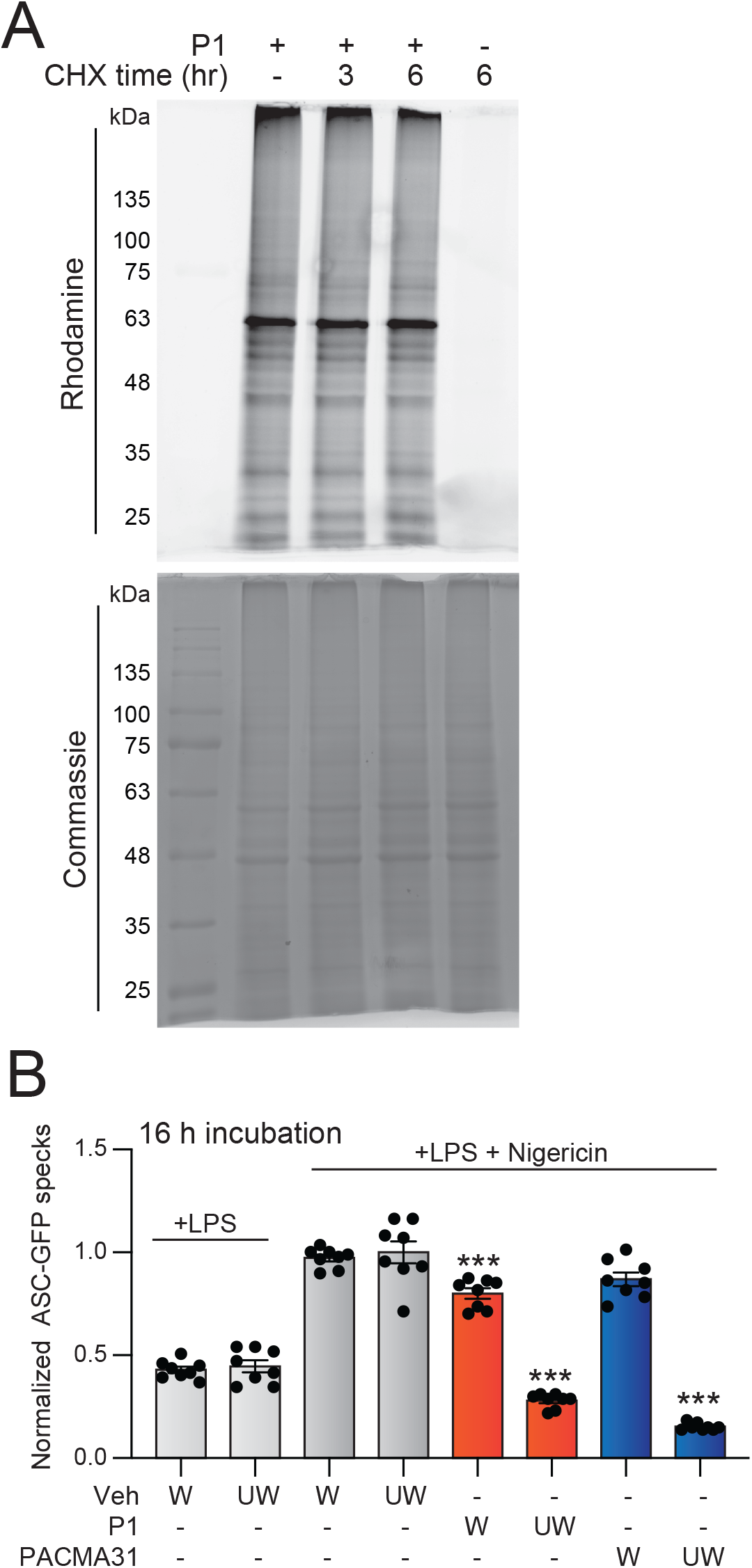
(Supplement to Figure 4) Acute treatment with P1 or PACMA31 blocks NLRP3 inflammasome activation independent of cellular toxicity. **A.** Fluorescent and Coomassie stained SDS-PAGE gels of lysates prepared from THP1 null cells primed for 16 h with LPS (1 µg/mL) and then treated with P1 (5 µM) for 1 h. Cells were then treated with cycloheximide (CHX; 50 µg/mL) for 0-6 h prior to lysis and click-dependent modification of the P1 conjugated proteins with rhodamine-azide. **B**. ASC-GFP speck formation from THP1 ASC-GFP pretreated for 16 h with LPS (1 µg/mL) and treated for 1 h with P1 (5 µM) or PACMA31 (5 µM). The cells were then washed, and the media was replaced with LPS in the absence (washed, W) or presence (unwashed, UW) of P1 or PACMA31. Nigericin (10 µM) was added for 2 h prior to imaging of ASC specks at the end of 16 h compound incubation. Error bars show SEM for n=8. ***p<0.005 for two-way ANOVA relative to vehicle-treated cells administered LPS and nigericin/ATP.

## REFERENCES

1 Sharma, B. R. & Kanneganti, T. D. NLRP3 inflammasome in cancer and metabolic diseases. Nat Immunol 22, 550–559, doi:10.1038/s41590-021-00886-5 (2021).

2 Swanson, K. V., Deng, M. & Ting, J. P. The NLRP3 inflammasome: molecular activation and regulation to therapeutics. Nat Rev Immunol 19, 477–489, doi:10.1038/s41577-019-0165-0 (2019).

3 Fu, J. & Wu, H. Structural Mechanisms of NLRP3 Inflammasome Assembly and Activation. Annu Rev Immunol 41, 301–316, doi:10.1146/annurev-immunol-081022-021207 (2023).

4 Xu, J. & Nunez, G. The NLRP3 inflammasome: activation and regulation. Trends Biochem Sci 48, 331–344, doi:10.1016/j.tibs.2022.10.002 (2023).

5 Aksentijevich, I. et al. The clinical continuum of cryopyrinopathies: novel CIAS1 mutations in North American patients and a new cryopyrin model. Arthritis Rheum 56, 1273–1285, doi:10.1002/art.22491 (2007).

6 Aganna, E. et al. Association of mutations in the NALP3/CIAS1/PYPAF1 gene with a broad phenotype including recurrent fever, cold sensitivity, sensorineural deafness, and AA amyloidosis. Arthritis Rheum 46, 2445–2452, doi:10.1002/art.10509 (2002).

7 Vandanmagsar, B. et al. The NLRP3 inflammasome instigates obesity-induced inflammation and insulin resistance. Nat Med 17, 179–188, doi:10.1038/nm.2279 (2011).

8 Yang, F. et al. NLRP3 deficiency ameliorates neurovascular damage in experimental ischemic stroke. J Cereb Blood Flow Metab 34, 660–667, doi:10.1038/jcbfm.2013.242 (2014).

9 Ding, S. et al. Modulatory Mechanisms of the NLRP3 Inflammasomes in Diabetes. Biomolecules 9, doi:10.3390/biom9120850 (2019).

10 Shao, B. Z. et al. Targeting NLRP3 Inflammasome in Inflammatory Bowel Disease: Putting out the Fire of Inflammation. Inflammation 42, 1147–1159, doi:10.1007/s10753-019-01008-y (2019).

11 Dinarello, C. A. Interleukin-1 in the pathogenesis and treatment of inflammatory diseases. Blood 117, 3720–3732, doi:10.1182/blood-2010-07-273417 (2011).

12 Brydges, S. D. et al. Inflammasome-mediated disease animal models reveal roles for innate but not adaptive immunity. Immunity 30, 875–887, doi:10.1016/j.immuni.2009.05.005 (2009).

13 Ismael, S., Zhao, L., Nasoohi, S. & Ishrat, T. Inhibition of the NLRP3-inflammasome as a potential approach for neuroprotection after stroke. Sci Rep 8, 5971, doi:10.1038/s41598-018-24350-x (2018).

14 Abbate, A. et al. Interleukin-1 and the Inflammasome as Therapeutic Targets in Cardiovascular Disease. Circ Res 126, 1260–1280, doi:10.1161/CIRCRESAHA.120.315937 (2020).

15 Mangan, M. S. J. et al. Targeting the NLRP3 inflammasome in inflammatory diseases. Nat Rev Drug Discov 17, 688, doi:10.1038/nrd.2018.149 (2018).

16 Yang, Y., Wang, H., Kouadir, M., Song, H. & Shi, F. Recent advances in the mechanisms of NLRP3 inflammasome activation and its inhibitors. Cell Death Dis 10, 128, doi:10.1038/s41419-019-1413-8 (2019).

17 Coll, R. C. et al. A small-molecule inhibitor of the NLRP3 inflammasome for the treatment of inflammatory diseases. Nat Med 21, 248–255, doi:10.1038/nm.3806 (2015).

18 Marchetti, C. et al. OLT1177, a beta-sulfonyl nitrile compound, safe in humans, inhibits the NLRP3 inflammasome and reverses the metabolic cost of inflammation. Proc Natl Acad Sci U S A 115, E1530–E1539, doi:10.1073/pnas.1716095115 (2018).

19 Coll, R. C. et al. MCC950 directly targets the NLRP3 ATP-hydrolysis motif for inflammasome inhibition. Nat Chem Biol 15, 556–559, doi:10.1038/s41589-019-0277-7 (2019).

20 Stanton, C. et al. Covalent Targeting As a Common Mechanism for Inhibiting NLRP3 Inflammasome Assembly. ACS Chem Biol 19, 254–265, doi:10.1021/acschembio.3c00330 (2024).

21 He, H. et al. Oridonin is a covalent NLRP3 inhibitor with strong anti-inflammasome activity. Nat Commun 9, 2550, doi:10.1038/s41467-018-04947-6 (2018).

22 Hooftman, A. et al. The Immunomodulatory Metabolite Itaconate Modifies NLRP3 and Inhibits Inflammasome Activation. Cell Metab 32, 468–478 e467, doi:10.1016/j.cmet.2020.07.016 (2020).

23 Bertinaria, M., Gastaldi, S., Marini, E. & Giorgis, M. Development of covalent NLRP3 inflammasome inhibitors: Chemistry and biological activity. Arch Biochem Biophys 670, 116–139, doi:10.1016/j.abb.2018.11.013 (2019).

24 Stanton, C. et al. The glycolytic metabolite methylglyoxal covalently inactivates the NLRP3 inflammasome. Cell Rep 43, 114688, doi:10.1016/j.celrep.2024.114688 (2024).

25 Rosarda, J. D., Stanton, C. R., Chen, E. B., Bollong, M. J. & Wiseman, R. L. Pharmacologic Targeting of PDIA1 Inhibits NLRP3 Inflammasome Assembly and Activation. Israel Journal of Chemistry 64, e202300125, doi:10.1002/ijch.202300125 (2024).

26 Medinas, D. B., Rozas, P. & Hetz, C. Critical roles of protein disulfide isomerases in balancing proteostasis in the nervous system. J Biol Chem 298, 102087, doi:10.1016/j.jbc.2022.102087 (2022).

27 Kline, G. M. et al. Divergent Proteome Reactivity Influences Arm-Selective Activation of the Unfolded Protein Response by Pharmacological Endoplasmic Reticulum Proteostasis Regulators. ACS Chem Biol 18, 1719–1729, doi:10.1021/acschembio.3c00042 (2023).

28 Paxman, R. et al. Pharmacologic ATF6 activating compounds are metabolically activated to selectively modify endoplasmic reticulum proteins. Elife 7, doi:10.7554/eLife.37168 (2018).

29 Hoffstrom, B. G. et al. Inhibitors of protein disulfide isomerase suppress apoptosis induced by misfolded proteins. Nat Chem Biol 6, 900–906, doi:10.1038/nchembio.467 (2010).

30 Ge, J. et al. Small molecule probe suitable for in situ profiling and inhibition of protein disulfide isomerase. ACS Chem Biol 8, 2577–2585, doi:10.1021/cb4002602 (2013).

31 Xu, S. et al. Discovery of an orally active small-molecule irreversible inhibitor of protein disulfide isomerase for ovarian cancer treatment. Proc Natl Acad Sci U S A 109, 16348–16353, doi:10.1073/pnas.1205226109 (2012).

32 Cole, K. S. et al. Characterization of an A-Site Selective Protein Disulfide Isomerase A1 Inhibitor. Biochemistry 57, 2035–2043, doi:10.1021/acs.biochem.8b00178 (2018).

33 Kline, G. M. et al. Metabolically activated proteostasis regulators that protect against erastin-induced ferroptosis. RSC Chem Biol 5, 866–876, doi:10.1039/d4cb00027g (2024).

